# Protein property prediction based on local environment by 3D equivariant convolutional neural networks

**DOI:** 10.1101/2024.02.07.579261

**Authors:** He Chen, Yifan Cheng, Jianqiang Dong, Jie Mao, Xin Wang, Yuan Gao, Yuchao Li, Chengzhi Wang, Qiong Wu

## Abstract

Predicting the properties of proteins is an important procedure in protein engineering. It determines the subspace of mutations for protein modifications, which is critical to the success of the project, but heavily relies on the knowledge and experience of scientists. In this study, we propose a novel deep 3D-CNN model, Eq3DCNN, specifically designed for local environment-related tasks in protein engineering. Eq3DCNN uses basic atom descriptors and their coordinates as inputs, utilizing customized data augmentations to enhance its training efficiency. To make the Eq3DCNN extracted features with more generalization capability, we incorporated a rotation equivariant module to get rotation invariant features. Using cross-validations with different data splitting strategies and under the scenarios of zero-shot predictions, we demonstrate that Eq3DCNN outperformed other 3D-CNN models in stability predictions, and also well-preformed on other prediction tasks, such as the binding pocket and the secondary structure predictions. Our results also identified the key factors that contribute to the model’s accuracy and the scope of its applications. These findings may help scientists in designing better mutation experiments and increasing the success rate in protein engineering.

## Introduction

Protein engineering is widely employed in both academia and industry to enhance the desired properties of proteins, such as modifying enzyme activities [1], improving antibody binding specificity [2], and increasing stability or yield for productions purposes [3, 4]. In practice, this process typically starts with a natural protein, followed by mutagenesis to generate a set of candidates (mutants), and screening these mutants to identify the ones with desired properties [5]. However, this strategy faces several challenges. First, it requires efforts and expertise to understand the structure and functional domain of the target protein [6]. Second, due to the tremendous mutational space, it is unrealistic to search the space entirely, requiring a critical design of the subspace of mutations [7]. Third, the evaluation of the engineered protein is laborious and expensive, usually with a high failure rate and requires repeating the process multiple times [8]. To address these challenges, computational methods that could understand the functional structure, explore a larger subspace of mutations, and efficiently evaluate the mutation result *in silico* are required.

Recently, deep learning methods have achieved remarkable success in many protein related tasks [7, 9, 10, 11, 12]. In particular, AlphaFold reached the state-of-the-art in protein folding prediction [13], and ESM demonstrated that protein sequence representations are generalized features for multiple applications [14, 15, 16]. However, these models mainly focused on the global or evolutionary features, which are more informative for their tasks. As for protein engineering, the local environment surrounding the mutation site is more relevant for predicting the properties of the engineered protein. Previous studies have demonstrated the utilization of a 3D convolutional neural network (3D-CNN) to extract local features from protein structures for such tasks [17]. However, one of the major challenges in using 3D-CNN is the computational complexity. The model size increases exponentially with the voxel size, resulting in high computational costs and requiring larger training datasets. The size is also related to the number of descriptors used as inputs, which could include the occupancy [18, 19], basic atom types [20, 21], biophysical or biochemical properties [22, 23], or other hand-crafted features [24, 25, 26, 27, 28]. In this way, recent reports usually employed shallow 3D-CNN models with a large set of hand-crafted features to achieve an acceptable model size. However, this approach limits the generalization capability of the 3D-CNN models and requires prior knowledge or assumptions for selecting the descriptors. Recalling to the 2D-CNN model used in computer vision [29], the current consensus is using the raw color pixels as input with deep models, which can extract abstract and generalized features from the images and outperform other methods. We believe this paradigm should also work in protein property predictions, and a deeper model with basic features as input could achieve better results if the model could be effectively trained.

Here, we proposed Eq3DCNN, a novel 3D-CNN model for local environment-related tasks in protein engineering, which employs coordinates of three basic atom types (carbon, nitrogen, and oxygen) as inputs. By introducing shortcut connections and rotation equivariance into the CNN block, Eq3DCNN could be effectively trained and extract features with more generalization capability. Compared with other 3D-CNN based methods, we evaluated our model and demonstrated its outperformance in thermostability predictions. Additionally, using a saturation mutagenesis dataset, we systematically analyzed the prediction results of Eq3DCNN and evaluated the factors that might affect the results. Furthermore, our model architecture exhibits promising generalization capacity to other tasks, such as the ligand-binding classification and the secondary structure predictions. These results suggested that Eq3DCNN has the potential to wide applications, and may increase the efficiency for protein engineering and drug discovery.

## Results

### The design of Eq3DCNN – a deep model with rotational invariance

To predict protein properties based on the local environment, we first need to define the center of interest (COI) and determine the size of the local environment in 3D space. We assume that the local environment around the COI contains sufficient information for the task. For example, when predicting mutational effects, the mutation site will be selected as the COI, and the size of the local environment will be a cube of 16 angstroms. Next, we perform voxelization to convert the local environment into a voxel tensor, which is a 3D volume of atom descriptors (with shape of 3 x 16 x 16 x 16, for example), with a resolution of 1 angstrom. Finally, the voxel tensor will be fed into Eq3DCNN, a deep 3D-CNN model, which extracts the key features of the local environment and makes predictions after several layers of nonlinear transformation (Figure 1A).

**Figure 1.**
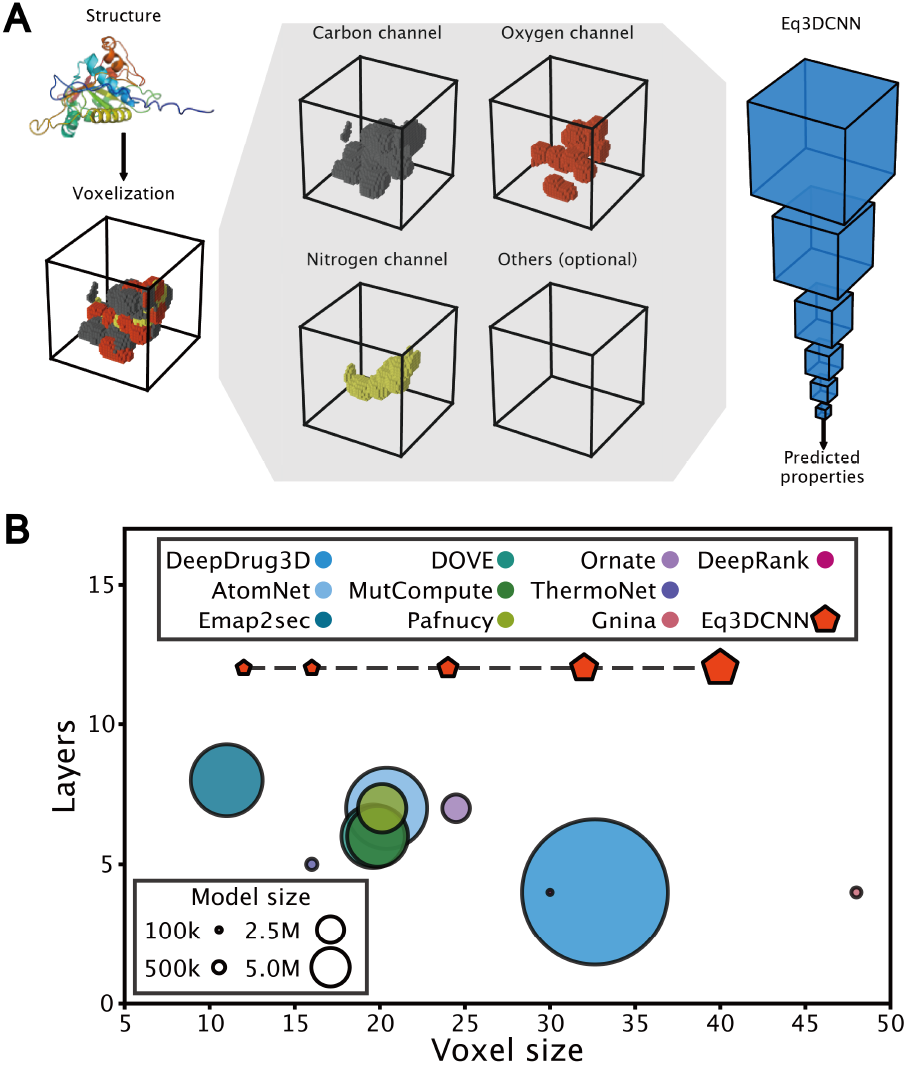
Overview of Eq3DCNN. (**A**) The workflow to predict local structure properties by Eq3DCNN. First, the center of interest (COI) and its surrounding local environment were extracted from the protein structure. Then, using three basic atom descriptors, the local environment was converted into a voxel tensor (voxelization). Last, the voxel tensor was fed into the Eq3DCNN and output the predicted properties. (**B**) Scatter plot shows the voxel size (x-axis), the number of layers (y-axis), and the number of model parameters (dot size) of Eq3DCNN and other 3D-CNN models.

Unlike 2D-CNN, training a deep 3D-CNN model is challenging in practice, especially in protein local properties prediction which only has a limited amount of data. Therefore, previous studies commonly used shallow 3D-CNNs with wide fully-connected layers (Figure 1B and Supplementary Table S1). Besides, the local environment of the protein and its corresponding properties are invariant to rotation in 3D space, which is more complex than in 2D space. Some previous studies used data augmentation to address this rotational invariance (Supplementary Table S1), which rotated the input voxel before feeding into the 3D-CNN model. However, this approach was unstable and inefficient for increasing data diversity due to voxel interpolation and edge effects – given a voxel (cube) of input features, only 24 rotations kept the data intact (6 upward faces × 4 rotations with 90°, referred to as intact rotations, Supplementary Figure S1A), and all other rotations (referred to as approximated rotations) break the integrity of the data (Figure 2A, Supplementary Figure S1B and C). To address these challenges, we modified the training pipeline and improved the model architecture by taking advantage of the invariance as an additional constraint (inductive bias). We first used heavy data augmentations to make the model more robust to small variations, especially at the voxel edges (Supplementary Figure S1D). This approach helps the model handle information loss and focus on key features in the local environment. Then, we applied a general equivariance framework to achieve rotational invariance [30, 31]. Finally, shortcut connections were added into the 3D-CNN block to make the model more robust and easier to train [32]. Combining these modifications, with increased data diversity and rotational invariance constraints, the Eq3DCNN could be efficiently trained and able to capture more generalized features.

**Figure 2.**
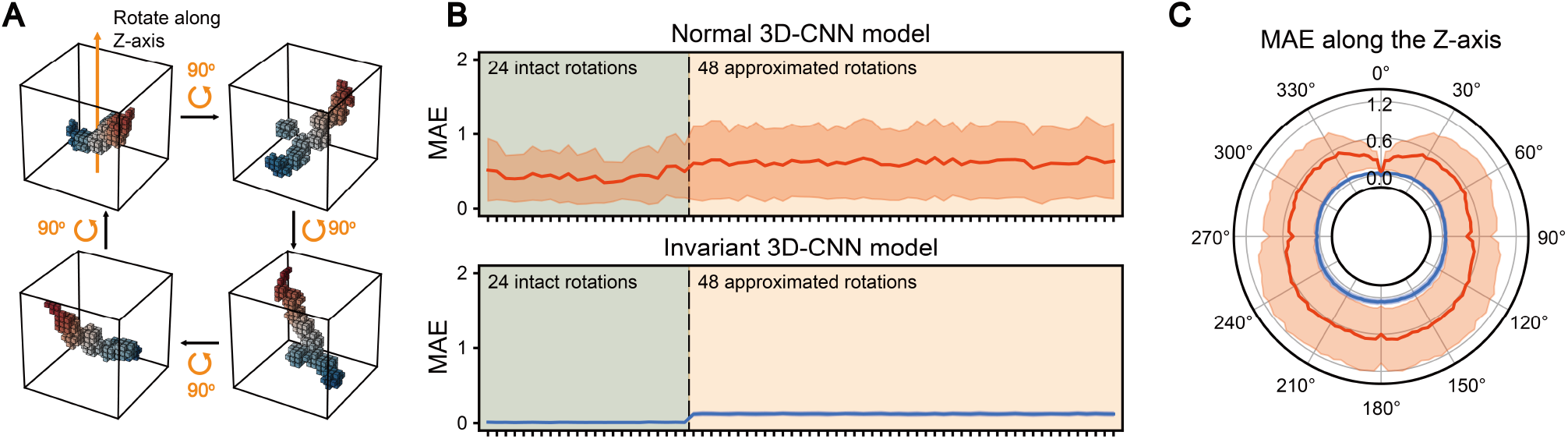
Rotational invariance of Eq3DCNN. (**A**) Demonstration of the protein local environment and rotational invariance. The voxel representation of a local environment was rotated along the z-axis four times of 90°. Despite the dramatic changes in the voxel representation, the local environment and its corresponding properties remain unchanged. (**B-C**) The mean absolute error (MAE) between outputs before and after rotated the input. The test was performed on a normal 3D-CNN (red) and Eq3DCNN (blue), respectively, using (**B**) 24 intact rotations and 48 approximated rotations or (**C**) continuous rotations along z-axis. Using different voxel representations (n=96) as inputs, the solid lines indicated the mean MAE of all outputs, and shade indicated the standard deviations.

To test the rotational invariance of Eq3DCNN, we used 24 intact rotations and 48 approximated rotations as inputs and examined the mean absolute error (MAE) between the outputs before and after the inputs were rotated. In a normal 3D-CNN model, the output changed dramatically after the input was rotated, and the approximated rotations exhibited larger MAE compared with the intact rotations (Figure 2B). Moreover, there were high variations among different inputs. In contrast, Eq3DCNN’s outputs with nearly zero MAE among all rotations, indicating its rotational invariance, even with the approximation rotations. To further check the invariance under other rotation angles, we completely tested a series of continuous rotations (0° to 360°) on a given axis (z-axis for example, Figure 2C). Compared with the original pose (0°), the MAE was constantly high in the normal 3D-CNN but negligible in Eq3DCNN. It was worth noticing that in normal 3D-CNN the error at 90°, 180°, and 270° were relatively lower because the data were intact under these rotations (Supplementary Figure S1C). But the error raised quickly even with a small degree of rotation, indicating the normal 3D-CNN was very sensitive to input changes. Taken together, with heavy data augmentations and shortcut connections, we successfully trained Eq3DCNN, a new deep 3D-CNN model that utilizes only three basic atom descriptors as inputs. Eq3DCNN could provide robust output with rotational invariance, even with missing data at the margin. This suggested that Eq3DCNN had the potential to extract more generalized features from the local environment, making it a better representation for many downstream applications.

### Benchmarking Eq3DCNN on thermostability prediction

To test the capability of Eq3DCNN, we applied the model on the point-mutation thermostability prediction task. We chose this task under the following considerations. First, this task is very practical in protein engineering since thermostability is an important property of proteins. Second, there is an amount of experimental data available about thermostability [33, 34, 35, 36], making it feasible to train a deep learning model. Third, the protein thermostability data are diversified, which makes it a complicated task that might reveal the power of the deep learning model, and also provide multiple ways to evaluate the model (discussed below).

We benchmarked our Eq3DCNN model and compared it with several other 3D-CNN models that have been reported in biological properties prediction (Supplementary Table S1). Two different strategies (Figure 3A) were used to separate the training and validation datasets: (1) randomly splitting the data (Rand-fold), which made the assessment generally predicting new mutants in the validation set, and (2) splitting the data by the protein (Prot-fold), which made the assessment focusing on predicting the “new” (unseen) proteins in the validation set. All the models were trained five times (k-fold cross-validation, k=5) under the same conditions and using the same input voxel size, then evaluated using the validation set (Figure 3B) and an extra testing dataset (Figure 3C). We tested the contribution of the three modification modules – data augmentation, rotation equivariant and shortcut connections - to the model performance improvement. The results showed that all of three could improve the performance, particularly the data augmentations and rotation equivariant modules, which could also be easily adapted to other models. Combining all three of our modifications, Eq3DCNN reached the highest correlation scores compared to other 3D-CNN models. It was worth noticing that even though some of these models were originally designed for other tasks (Supplementary Table S1), they still showed acceptable performance in this thermostability prediction task. Comparing the two splitting strategies (Rand-fold and Prot-fold) on the validation set, the correlations between the predicted and ground truth values were lower when using the Prot-fold, and the variations among folds were also higher when using the Prot-fold than using the Rand-fold. These results suggested that predicting the properties for unseen proteins was more challenging.

**Figure 3.**
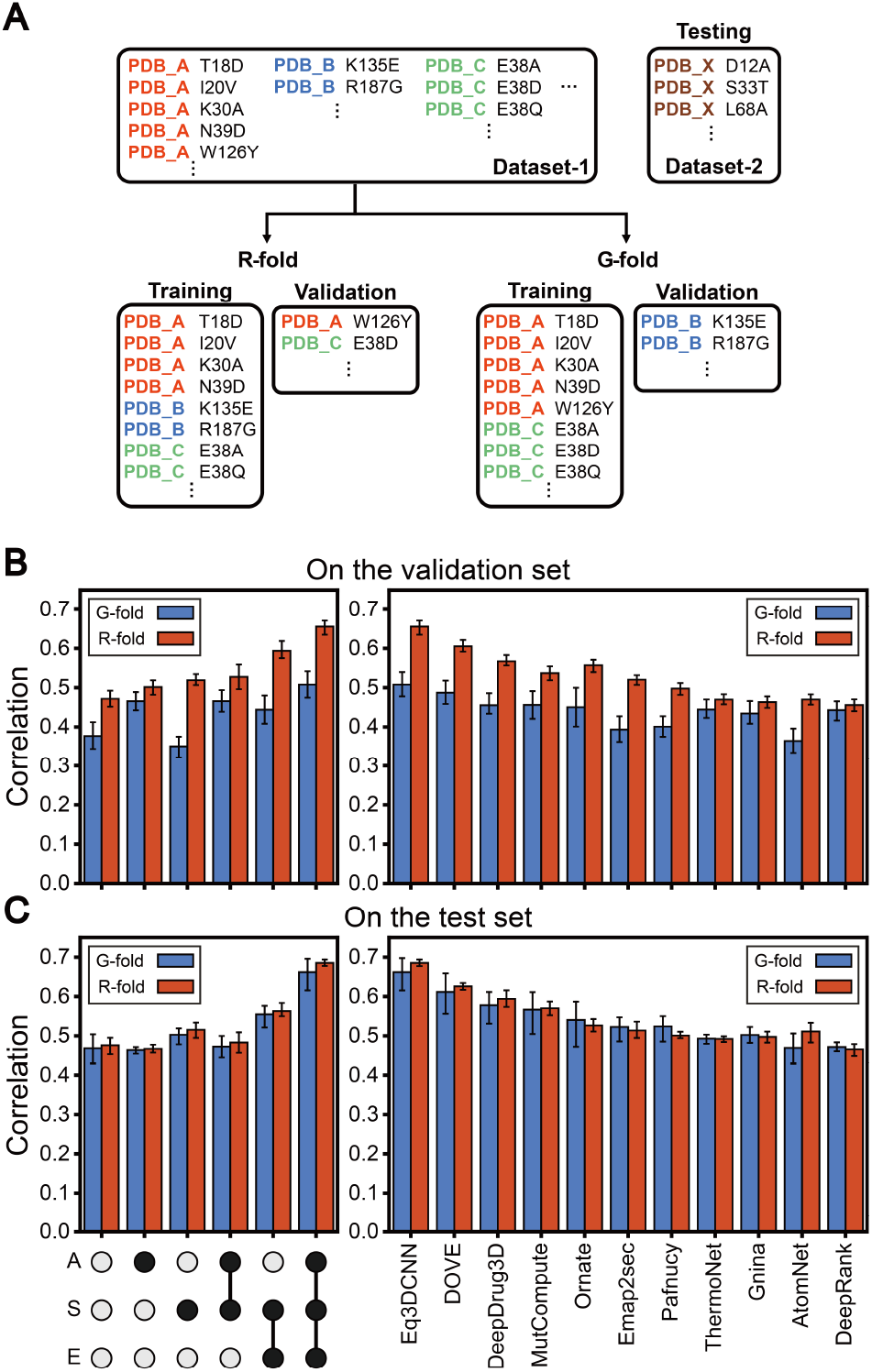
Benchmarking Eq3DCNN on thermostability prediction task. (**A**) The setup of benchmarking datasets. Two datasets were used for training, validation, and testing. Two splitting strategies were used to evaluate the prediction of new mutations (Rand-fold) and new proteins (Prot-fold) for each dataset. (**B-C**) The benchmarking results in (**B**) validation set and (**C**) testing set. Barplot shows the Spearman rank correlation between the predicted and ground truth values. The left panel: barplot benchmarking with the corresponding selections of the three of modification modules. (A for data augmentations, S for shortcut connections, and E for rotation equivariant). The right panel: barplot benchmarking on Eq3DCNN and other 3D-CNN models. The error bar indicated the standard deviations among 5-folds.

### Testing Eq3DCNN on synthetic proteins

To investigate the generalization capability of Eq3DCNN, we tested the model on a site-saturation mutagenesis dataset (referred to as SSM2) of synthetic proteins to predict the protease stability changes (Δ*S*) among mutations [37]. Considering the positive correlation between protease stability and thermostability [37], we first treated it as a zero-shot prediction task, in that we trained the model using the previous thermostability dataset and inferred on the synthetic protein dataset. Despite the synthetic proteins were not found in nature, the zero-shot predicted values showed positive correlation with the protease stability values, and Eq3DCNN outperformed other 3D-CNN models (Supplementary Figure S2A). However, the correlation score from Eq3DCNN results was about 0.35, which was significantly lower than that on thermostability dataset (0.50-0.66, Figure 3B). It was also lower than the reported correlation between thermostability and protease stability (0.7-0.8) [37], indicating that the difference between the two datasets was non-negligible. Meanwhile, researchers usually focus on the mutants with relatively better or worse properties rather than the full mutational space, making the correlation score less meaningful and non-intuitive in practice. Hence, we proposed a new rank-based evaluation method by simulating a protein optimization process. Specifically, for each protein, we sorted all the mutants based on their predicted and ground truth Δ*S* respectively (Supplementary Figure S2B). Then, under a given target percentage (for the highest or lowest percentage of the mutational space), we calculated how many of them were overlapped, which we refer to as the success rate (Supplementary Figure S2C). For example, for a given protein with 1000 mutants, if we aim to get the top 100 unstable (or stable) mutants (the target percentage is 10%), based on our result, about 14-44 of the top 100 mutants predicted by the model were reliable (the success rate ranging from 14-44%, varying among different proteins). Compared with the result of random sort, the Eq3DCNN significantly outperformed when the target percentage is low (Figure 4A and Supplementary Figure S2D). As the target percentage was raised, the success rate increased, and the difference between Eq3DCNN and random sort became similar. This rank-based evaluation method provided informative and instructive results, which indicated the applicability of the model and suggested the sample size for the wet-lab experiment. However, a very high criterion screening is not effective using this method. If the goal was to get the highest 1% mutants, the success rate was very low (less than 1%), almost equal to the random results. Furthermore, under a given target percentage, the sample size was correlated with the size of the mutational space, and the success rate was varying among different proteins. When applying such methods in practice, researchers must compromise among the target percentage, the success rate, and the sample size to maximize the effectiveness of the method.

**Figure 4.**
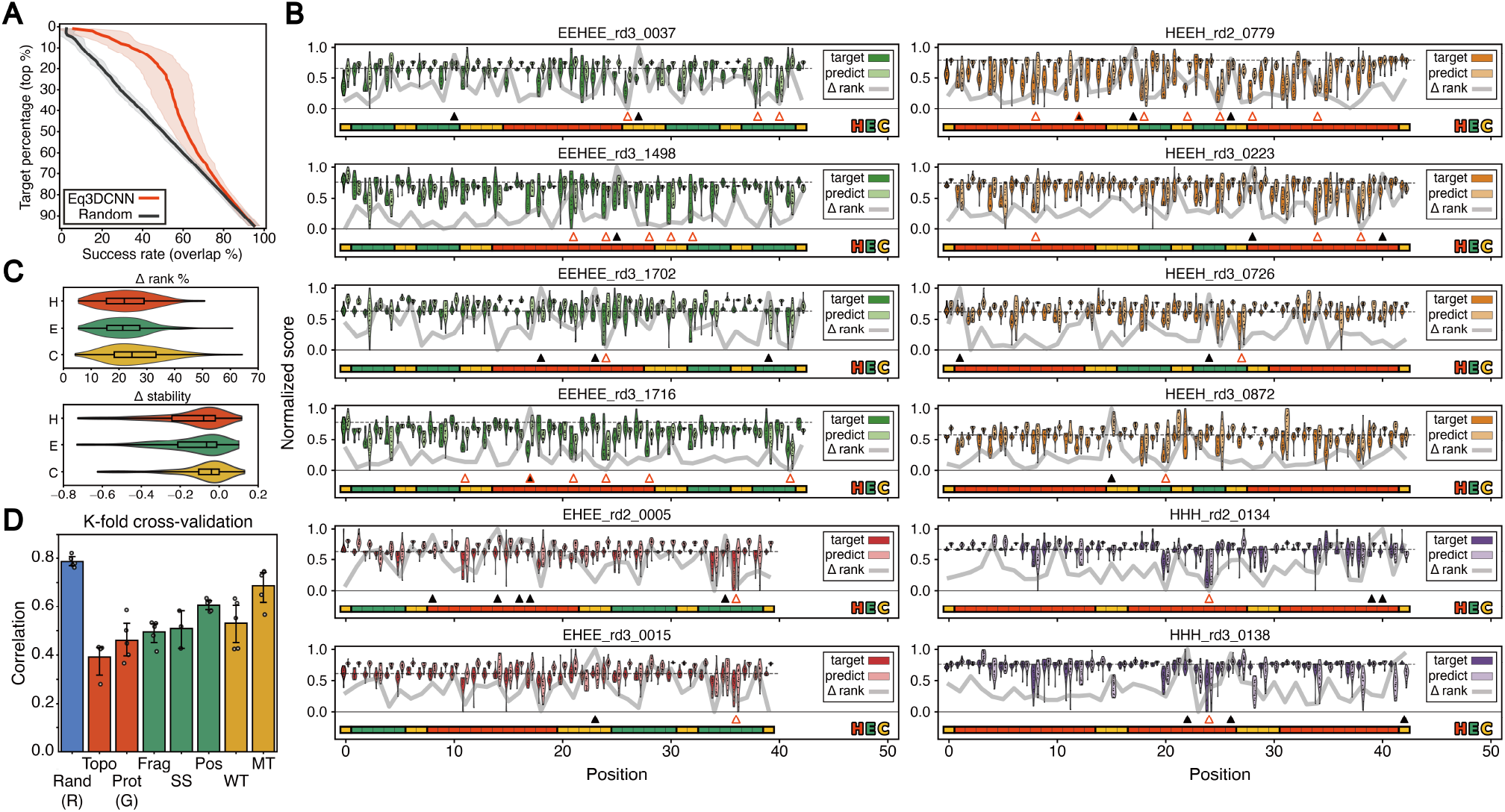
Zero-shot and analysis on stability prediction of synthetic proteins. (**A**) A rank-based evaluation result by simulating the protein optimization process. For a given target percentage (y-axis), the success rate (x-axis) was the overlapped percentage between the prediction and ground truth results. The solid lines indicate the Eq3DCNN (red) result and random (black) result. The shade indicates the standard deviation among different proteins. (**B**) The landscape of the zero-shot predicted and ground truth Δ*S* (normalized, y-axis) for each position (x-axis). Each panel shows the results of a protein in the dataset, and the violin plot with dots indicates the distribution of predicted and ground truth Δ*S*. The dashed line indicates the values of the wildtype, and the thick gray line indicates the prediction accuracy (mean of Δrank for each position, the lower the better) about the corresponding position. The underlying colorbar indicates the secondary structure of the protein. H (red) for *α*-helix, E (green) for *β*-strand, and C (yellow) for coil. The crucial positions (red triangle) and failure positions (black triangle) are labeled. (**C**) Violinplots indicate the distribution of Δ*S* and Δrank% on three secondary structure. (**D)** Barplot shows the Spearman rank correlation using 8 different k-fold separation strategies.

To get more details about the prediction results, we further investigated the distribution of the predicted and ground truth Δ*S* across different proteins and mutation positions (Figure 4B). In order to represent the prediction accuracy for each position, we first tried using the binary classification accuracy (stable Δ*S* > 0 or unstable Δ*S <* 0) or MAE to represent each position (Supplementary Figure S3). However, both of them turned out to be inappropriate for evaluation. The former approach overrepresented the mutations with slight effects around the wildtype (Δ*S* ≈ 0), while the latter approach exhibited a bias to the stability changes with larger values (such as |Δ*S*| > 1). Therefore, we used the rank difference between the predicted and ground truth Δ*S* for each protein as a score (Δrank, the lower the better) to indicate the prediction accuracy. Then, we used the mean Δrank of all mutations at the same position to represent the accuracy of the position. The results showed that the predicted Δ*S* aligned well with the ground truth Δ*S* at most of the positions, without obvious bias to any specific protein or positions (Figure 4B). However, we identified some positions where the prediction values were inconsistent or even opposite to the ground truth Δ*S* (failure positions, labeled by black solid triangle). Considering the distribution of ground truth Δ*S*, most positions only had minor effects on stability, with a few positions changed the stability dramatically (crucial positions, labeled by red triangle). Compared with other positions, our model performed well on such crucial positions (Supplementary Figure S2D and Figure S4), probably because the crucial positions with significant differences were easily captured by the model. This feature has potential benefits for various protein engineering applications. Furthermore, we investigated the impact of the secondary structure on the mutation position, to check whether it affects the prediction accuracy (Figure 4C). The Δrank on the coil regions (labeled by C) was slightly higher than that on the *α*-helix regions (labeled by H) and *β*-strand regions (labeled by E). This might be caused by the distribution of Δ*S*, which was close to zero on the coil, making it harder to distinguish. Taken together, after a comprehensive analysis of the Eq3DCNN results of zero-shot prediction, we demonstrated that there was no systematic bias present in the prediction results.

To further investigate other factors that might affect the prediction accuracy, we trained Eq3DCNN on the SSM2 dataset and evaluated it using the k-fold cross-validation results with different splitting strategies, similarly with what we had used previously in Figure 3. We tested eight different k-fold splitting strategies (Figure 4D and Supplementary Figure S5), which will be discussed in more detail below. We first analyzed the results of Rand-fold and Prot-fold, and compared the correlation score of the SSM2 and the thermostability dataset. It turned out that the correlation score of Rand-fold was higher on the SSM2 dataset compared to the thermostability dataset (0.79 vs 0.66), while the correlation score of Prot-fold was slightly lower (0.46 vs 0.50). This was probably due to the differences between the dataset composition, where the SSM2 dataset is a site-saturation mutagenesis dataset with only twelve proteins, each with over 710 mutants, whereas the thermostability dataset contains hundreds of proteins with an average of 21 mutants each. As a result, the model trained on the SSM2 dataset performed better on mutants (Rand-fold), while the model trained on the thermostability dataset had better performance on proteins (Prot-fold). When comparing the results on the SSM2 dataset, the difference in correlation results reflected the difference in dataset composition, which was only introduced by the splitting strategies. For example, splitting by the protein topology (Topo-fold) was a set of special case of Prot-fold (Supplementary Figure S5A and B), which incorporated the topology information that further reduced the diversity of the training data, resulting in the worst performance. When splitting dataset by positions (Supplementary Figure S5C), our hypothesis was that the random position splitting (Pos-fold) would have the best performance, followed by the Frag-fold, which split the dataset by a continuous fragment of positions, and the SS-fold, which incorporated the secondary structure information, with the worst performance. However, we found that the SS-fold and Frag-fold had similar correlation scores, indicating that secondary structure may not play a critical role for the prediction accuracy. Furthermore, we conducted a comparison of the dataset splitting methods based on the amino acid. Specifically, the WT-fold that splitting the dataset by the wildtype amino acid, while the MT-fold that splitting the dataset by the mutation amino acid. Since wildtype amino acids are associated with specific positions, the WT-fold could be considered as a special case of Pos-fold that incorporates the amino acid information. As a result, the correlation score of the WT-fold was lower than that of the Pos-fold. On the other hand, due to the nearly identical input voxels for different mutations at the same position, the MT-fold had more similar distributions of input between training and validation datasets, resulting in high correlation score. Overall, our analysis suggested that the protein type (Topo-fold, Prot-fold) was the most important factor affecting the prediction accuracy. The fragment of positions (Frag-fold, SS-fold) and the wildtype amino acids (WT-fold) had intermediate importance, while the positions (Pos-fold) and mutation amino acids (MT-fold) were the least important factors. These valuable results could help researchers to design the mutation strategy or subspace.

### Applying Eq3DCNN to other four prediction tasks

Besides stability, there are other properties that mainly rely on the local environment of proteins. To test the performance of Eq3DCNN in predicting these properties, we designed four different tasks to cover a wide range of applications. The first task was the protein-ligand affinity prediction (binding regression) [26, 27, 38], a regression task to predict the affinity (*K*_*d*_) for a given protein-ligand pair, in which the center of interest (COI) was set to be the center of the binding pocket. The second task was the protein pocket classification [21, 28], a binary classification task to determine whether a given position was within a binding pocket, in which the COI was set to be that position. These two tasks were related to the protein-ligand virtual screening and docking, which could help accelerate drug discovery and development. For these two tasks, the voxel size might significantly impact the results. Hence, we tested Eq3DCNN using different voxel sizes. The results showed that as the size increased, the accuracy improved at first. But after the voxel size was bigger than 24 angstroms, the accuracy either dropped (Figure 5A) or changed negligibly (Figure 5B). However, the computational costs increased exponentially (Figure 5C). These results suggested that the voxel size ranging from 16 to 24 angstroms was appropriate for these applications. We also examined the effect of different atom descriptors on the prediction results, by comparing the results of basic and complex atom descriptors (Figure 5D, E). As we expected, using different atom descriptors barely changed the prediction accuracy.

**Figure 5.**
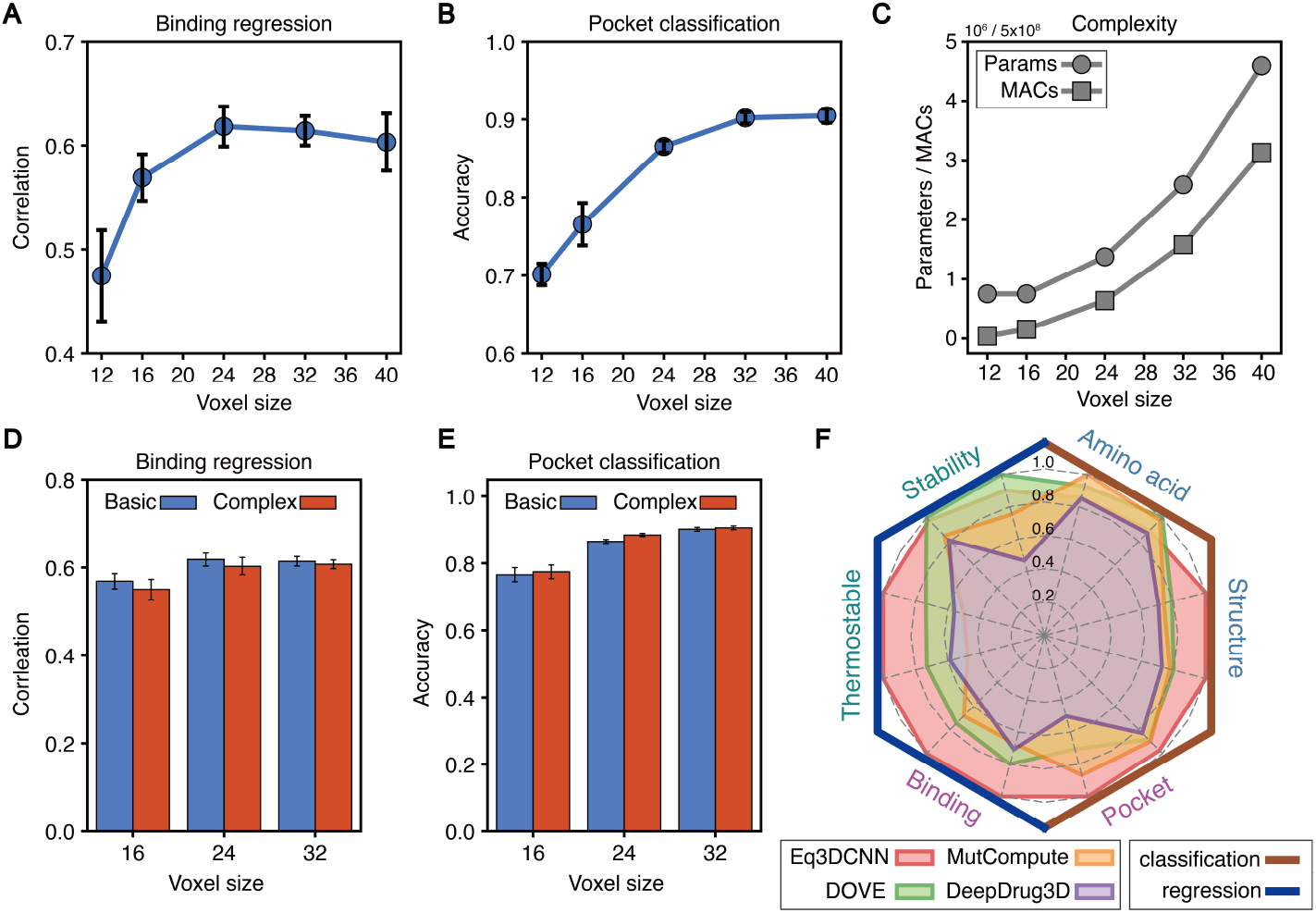
Testing Eq3DCNN on ligand-binding and self-supervised prediction tasks. (**A**) Voxel size and prediction accuracy for the binding affinity regression task. The accuracy was represented by the Spearman rank correlation. (**B**) Voxel size and prediction accuracy for the pocket classification task. (**C**) Voxel size and computational costs, which were represented by the model size (number of trainable parameters) and multiply-accumulate operations (MACs) (**D** and **E**) Comparing the results of Eq3DCNN between using basic descriptors and complex descriptors on (**D**) binding affinity regression task and (**E**) pocket classification task. (**F**) The radar plot shows the overall performance on six different tasks covering two types (classification and regression). The accuracy scores were min-max normalized with eleven 3D-CNN models. For each task, the two results under different training setups were shown.

The third and fourth tasks were to predict the secondary structure [19] and to predict the amino acid [24] for a given position, respectively. The COI was set to be the position of the amino acid and the center of the voxel tensor was masked to remove the target information. These two tasks could be used in structure biology [19, 22] and synthetic biology [24, 39, 40] researches. Additionally, for these two tasks, the target labels were generated from the input data, known as a self-supervised learning, which could be used in developing new models for few-shot or zero-shot learning to deal with other tasks that only have rare data.

To summarize all the results of this work, we classified them into six categories of tasks (thermostable, stability, amino acid, structure, pocket, and binding), covering two types of predictions (classification and regression) (Figure 5F). For each task category, the results under two different training setups were used, and the prediction scores (accuracy for classification tasks, Spearman correlation for regression tasks) were min-max normalized among all models to represent the prediction accuracy. Compared with the other three models with top performance, Eq3DCNN exceeds almost all of them, indicating a better generalization potential than other 3D-CNN based methods.

## Discussion

In this work, we developed a new method named Eq3DCNN, which predicts protein properties based on the local environment on the 3D structure. By leveraging a rotation equivariant module and implementing customized data augmentations, Eq3DCNN outperformed other 3D-CNN methods in several prediction tasks, including the stability prediction and the ligand-binding prediction. The most effective change of Eq3DCNN was the rotation equivariant module, which was implemented via group convolution of finite group [30]. This approach is easily adapted to other 3D-CNN models, and comparable to normal 3D-CNN without changing the architecture. Theoretically, the steerable convolution may achieve better accuracy with more generalized features [41, 42], which is also less computationally expensive and able to handle infinite groups. Unfortunately, we did not find a valid set of hyperparameters for the steerable convolution model. There is also a lack of reference using steerable convolution in the field of protein engineering, which still requires further investigation.

In the most of real-world cases, the target protein often lacks experimental data as a reference. Therefore, we systematically tested and analyzed the prediction results of Eq3DCNN under the zero-shot prediction scenario, using a site-saturation mutagenesis dataset of synthetic proteins [37]. The results of Eq3DCNN were satisfactory, with no significant bias on position or secondary structure. We also demonstrated the limitation of the methods under the view of protein engineering, as shown in Figure 4A. In the absence of experimental data, it is unrealistic to identify the mutant with best property. However, the Eq3DCNN could improve the success rate by obtaining a set of mutants with better chance of desired property. It is important to note that for some proteins with many functional studies, the starting point for bioengineering was already through a series of optimizations by biochemists. This optimization process could raise the actual target percentage, and result in a decrease of success rate for modifications. One potential solution to address this problem is incorporating the experimental data into the training dataset, which could be newly generated data or collected from previous studies. Overall, estimating the target percentage and determining the strategy still requires some experience and knowledge about the target protein.

To our knowledge, we are the first to employ the k-fold cross-validation with different groups to investigate the key factors that affect the prediction accuracy (Figure 4C). Our findings indicate that the protein type (topology and overall structure) has the greatest impact, followed by local structural attributes such as amino acid types. These results provide valuable insights into the capabilities and limitations of 3D-CNN based methods. This finding could help the researchers to design the subspace of mutations, and to maximize the power of the model and the experimental data. For example, in the worst case where the target protein and its fragments have no homolog in the dataset, the prediction accuracy will be relative lower than the protein with homologs. To improve this, one potential solution is the two-stage strategy. At stage one, scientists could use the model under the generalization (Prot-fold or Topo-fold) or zero-shot scenario, then select a collection of candidates with widespreading positions and preform the experiment. At stage two, the experiment results can be incorporated into the training dataset, to reach a higher accuracy (Pos-fold or Rand-fold). We believe this wet-lab and dry-lab combined strategy is the best solution for protein engineering.

Furthermore, the Eq3DCNN were tested in two completely different scenarios: the thermostability regression with the mutation site as the COI, and the pocket classification with the local environment center as the COI. For a fair comparison with other 3D-CNN based methods, we trained all the models under the same conditions. The Eq3DCNN outperformed other models in both cases, suggesting it has the potential of broad applications in protein engineering. However, training a model with such general capability typically requires a large amount of data that cover as many contexts as possible. In protein engineering, the amount of labeled data is very limited, or often restricted to a few proteins relying on high-throughput experiments [4, 43, 44, 45]. As demonstrated by our cross-validation analysis, the protein type is the most important factor, implying that data from previous high-throughput experiments are insufficient for training a model with general applicability. On the other hand, the unlabeled data is relatively abundant, both in quantity and diversity. Therefore, we believe using self-supervised learning strategy is a more promising approach to train a generalized model. In this manuscript, we also demonstrated applying self-supervised learning strategy on 3D-CNN using two different proxy tasks, the amino acid prediction and the secondary structure prediction. The performance and potential applications of these pre-training models are still worth further investigation.

## Materials and Methods

### Eq3DCNN and input features

The Eq3DCNN model was composed of two components: a 3D-CNN module and a fully-connected module. The 3D-CNN module had one 3D-CNN layer and four ResNet-like 3D-CNN blocks. The first 3D-CNN layer was designed with a kernel size of 5, and 16 output channels. For the four ResNet-like 3D-CNN blocks, each block had two 3D-CNN layers with a kernel size of 3 and a shortcut path. Batch normalization was employed. The output channels of the four blocks were 16, 32, 64, 128, respectively. Down-sampling were applied in the first two blocks and the last block. The fully-connected module had two linear layers with 256 and 64 channels respectively, and a final layer with variable number of channels, depending on the task. The ReLU activation were used for all the layers. The escnn package [46] was used to build the rotation equivariant 3D-CNN and get the invariant features. The model was implemented using PyTorch (v1.13).

For the input features, we used three basic atom descriptors (the carbon, nitrogen, and oxygen atoms). We also tested the complex atom descriptors, which used seven biophysical or biochemical properties (hydrophobic, aromatic, hydrogen-bond acceptor, hydrogen-bond donor, positive ionizable, negative ionizable, and occupancies). The atom descriptors and voxelization were extracted using the MoleculeKit package [47].

### Training and data augmentations

Eq3DCNN and other models were trained using the Adam optimizer with an initial learning rate of 0.001, accompanied by cosine annealing scheduler. The smooth L1 loss was used for the regression tasks, while the cross-entropy loss was used for the classification tasks.

In training, we used a series of data augmentations: (1) Random Rot: rotate the data along a random axis by an arbitrary angle or random choice one of the 24 intact rotations; (2) Random Noise: random noise (Gaussian noise) was introduced into the data; (3) Random Dropout: randomly selected some fraction of the data and set to zero; (4) Random Flip: randomly selected one axis (x, y, or z) and flipped the data along that axis. This mirror reflection transformation breaks the chirality of the data, but we considered the chirality was irrelevant for the prediction tasks; (5) Random Roll, random select one axis (x, y, or z) and roll the data along that axis. Some of the data was moved out and moved into the voxel cube from another boundary, which required the Center Crop [Padding] to remove this artifact; (6) Center Crop [Padding], crop a small cube of the data at the center and remove (set to zero) the surroundings. (7) Center Crop [Masking], remove (set to zero) a small cube at the center of the voxel. This was only used in self-supervised tasks (amino acid and the secondary structure predictions) to remove the target information.

These data augmentation methods were directly applied to the voxel features instead of the protein structure files (PDB), which could be effectively applied to the training pipeline, but might raise the information loss at the edges. These information loss and approximation were also considered as an additional step of data augmentation.

### Datasets and prediction tasks

#### Thermostability and stability regression tasks

For these two tasks, we constructed the dataset according to the previous report [23]. Specifically, the tasks were to predict the thermostability or stability changes between wildtype (WT) and mutant with a single point mutation (MT). The center of interest (COI) was set to be the position of the mutation, and the voxel size was 16. The concatenated voxels of wildtype and mutant were used as the input, and the Pearson correlation and Spearman correlation were used for evaluation. For the thermostability prediction task, the Gibbs free energy of folding Δ*G* was used, and the target was ΔΔ*G* = Δ*G*_WT_ −Δ*G*_MT_. The Q6428 dataset [23] was used for training and validation, which had 148 unique proteins (PDB IDs) with 6,428 records. The S^sym^ dataset [23] was used for testing, which had 15 unique proteins with 342 records. Both datasets were collected from the ProTherm database [33]. For the stability task, the stability score (*S*) was used, and the target was Δ*S* = *S*_WT_ − *S*_MT_. The dataset was extracted from previous study [37], and only the *de novo* designed miniproteins of four topologies were kept, which had 12 proteins with 9106 records in total.

#### Binding regression and pocket classification tasks

For these two tasks, the raw data and metadata were downloaded from PDBbind (the refined set, version 2020) [38], which had 5316 records. For the binding regression task, the structure of the binding pocket was used. The COI was set to be the center of the binding pocket, and the − log *K*_*d*_ was used as the target, with the Spearman correlation for evaluation. For the pocket classification task, the structure of the whole protein was used. For each protein, we randomly sampled more than 4 positions within the pocket as the positive samples, and same number of positions outside the pocket as the negative samples, which resulting in a balanced dataset. The COI was set to be the position, and the binary classification accuracy was used for evaluation. For these two tasks, five different voxel sizes (12, 16, 24, 32, 40) were tested. Some records failed in the voxelization, after removing those failure records, we obtained 5,214 records of unique proteins for the binding regression task, and 2,592 proteins with 19,766 records for the pocket classification task.

#### Self-supervised classification tasks

For these two tasks, the raw data and metadata were downloaded according to the previous report [25], which had 17k proteins with 1,456k records. The original dataset covered at most 100 or 50% of the residues for each protein, compromised with the computational costs, we downsampled the dataset 20 times, and obtained ∼15k proteins with ∼79k records. To remove the target information and constructed the self-supervised task, the center of the 6 angstroms cube was removed after voxelization, using the “Center Crop [Masking]” method.

## Author contributions statement

H.C. designed and conducted the research, H.C. and Y.C. performed the experiments. H.C., X.W., Y.G. and Y.L. designed the tasks, H.C. and Y.C. prepared the data, H.C. and Y.C. wrote the code, Y.C. trained the models, H.C. and Y.C. analyzed the results. H.C., Y.C., J.D., J.M., X.W., Y.G., Y.L., C.W., and Q.W. wrote and reviewed the manuscript, C.W. and Q.W. supervised the research.

## Acknowledgments

The authors thank the other group members for helpful discussion. The authors received no funding support for this research.

---

**Supplementary Table S1.** All 3D-CNN models tested in this study.

| Model | Task | Type | Voxel | Layers<sup>1</sup> | Augmentation<sup>2</sup> | Reference |
| --- | --- | --- | --- | --- | --- | --- |
| AtomNet | ligand-binding | classification | 20 | 7 | no | [18] |
| Emap2sec | secondary structure | classification | 11 | 8 | no | [19] |
| DeepDrug3D | ligand-binding | classification | 32 | 4 | no | [20] |
| DOVE | docking | classification | 20 | 6 | yes | [21] |
| Ornate | structure quality | regression | 24 | 7 | no | [22] |
| ThermoNet | stability | regression | 16 | 5 | yes | [23] |
| MutCompute | amino acid | classification | 20 | 6 | no | [24, 25] |
| Pafnucy | ligand-binding | regression | 20 | 7 | yes | [26] |
| Gnina | ligand-binding | classification / regression | 48 | 4 | yes | [27] |
| DeepRank | docking | classification / regression | 30 | 4 | yes | [28] |

<sup>1</sup>Number of 3D-CNN and linear layers.<sup>2</sup>Rotation-based data augmentation, not including symmetry-based data augmentation.

**Supplementary Figure S1.**
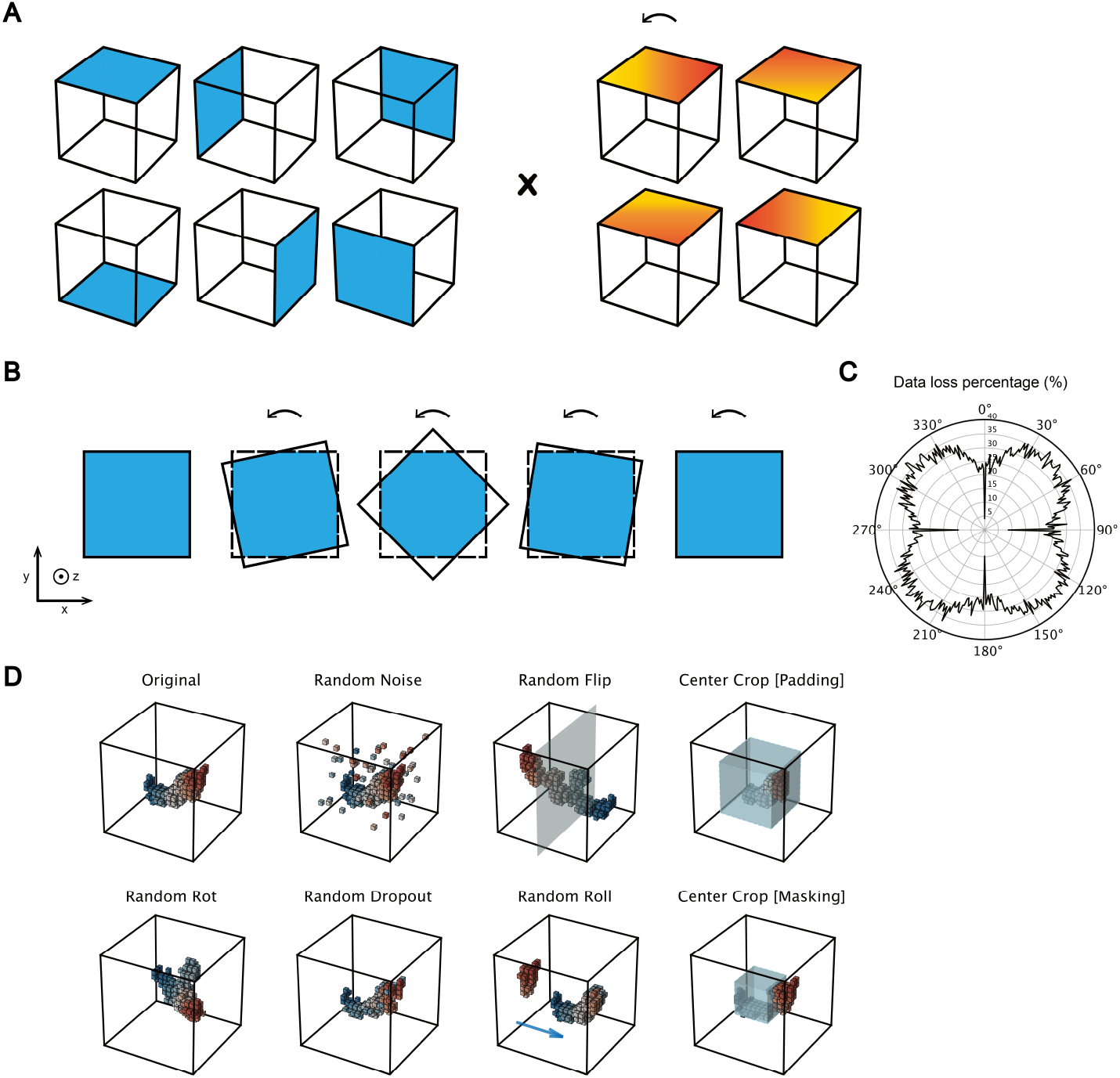
Demonstration of rotation and data augmentation. (**A**) Demonstration of the 24 intact rotations, 6 upward faces × 4 rotations with 90°. (**B**) Demonstration of rotations with information loss. (**C**) Example of information loss under continuous rotations, using simulated data. Such information loss was not only due to rotation but also as a result of approximation in voxelization. (**D)** Demonstration of the process of data augmentations.

**Supplementary Figure S2.**
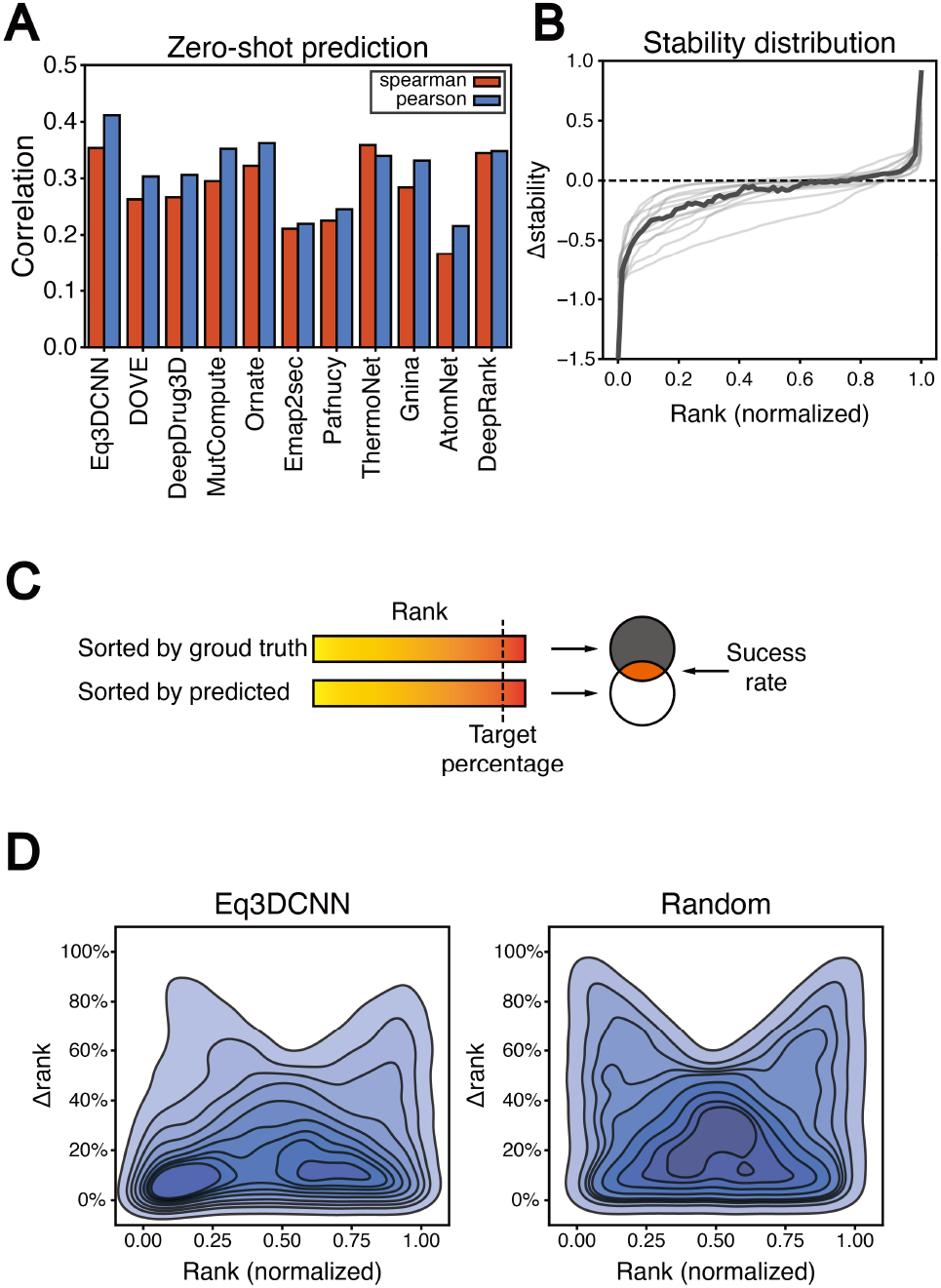
Zero-shot prediction results on SSM2 and rank-based evaluation. (**A**) Zero-shot prediction results in SSM2 dataset. (**B**) The distribution of stability scores and ranks in SSM2 dataset. Most of the mutants with slight effects around the wildtype (Δ*S* ≈ 0). (**C**) Demonstration of the rank-based evaluation method. (**D)** The distribution of rank and Δrank of Eq3DCNN and random results.

**Supplementary Figure S3.**
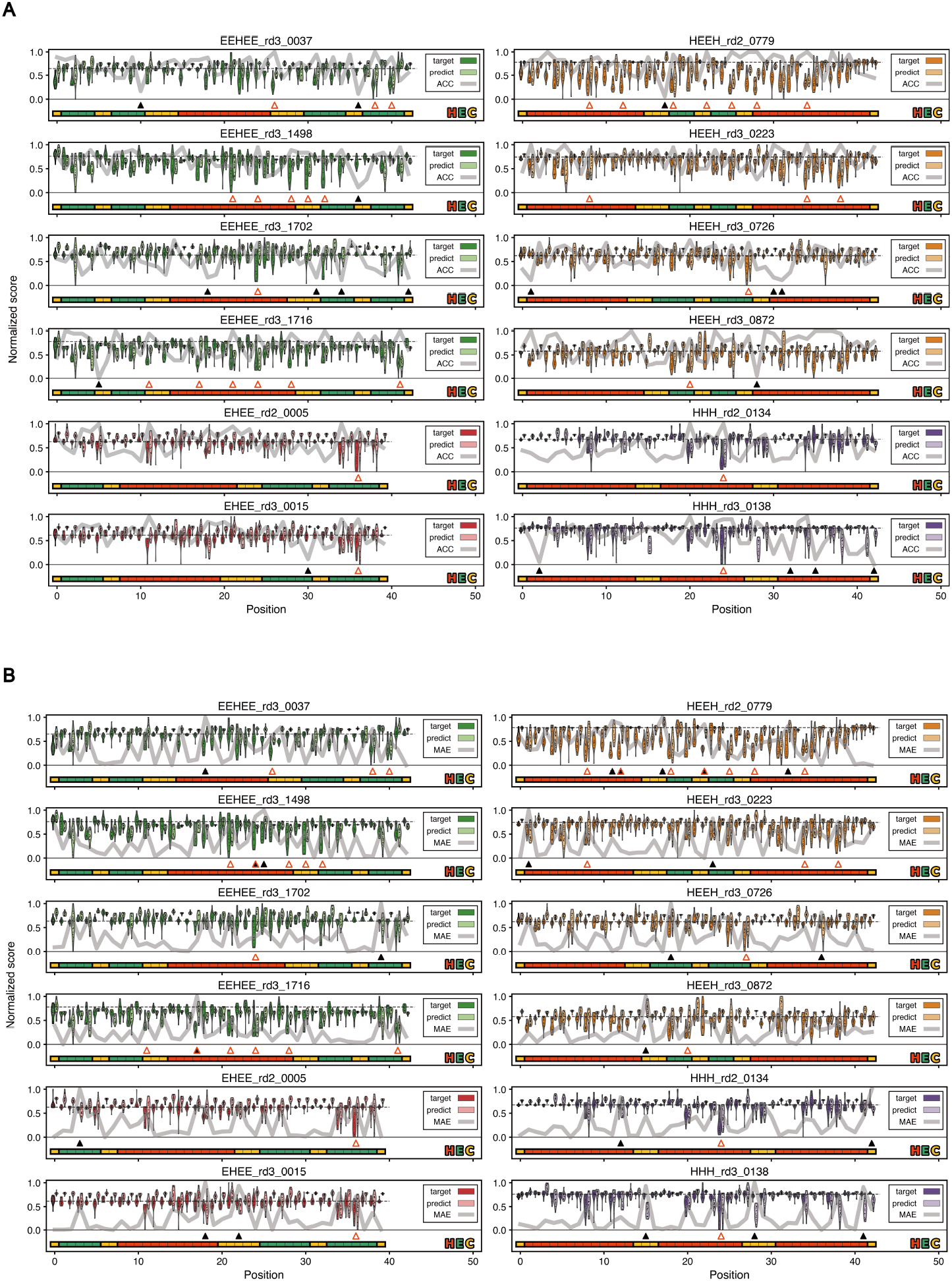
The landscape of the zero-shot predicted Δ*S* and ground truth Δ*S*. The prediction accuracy for each position represented by (**A**) binary classification accuracy (ACC) and (**B**) MAE.

**Supplementary Figure S4.**
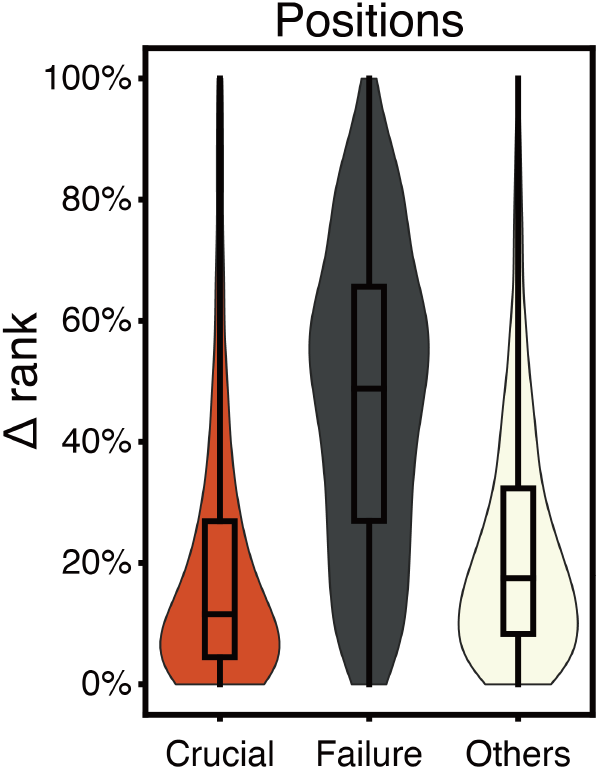
Distribution of prediction accuracy on crucial, failure and other positions.

**Supplementary Figure S5.**
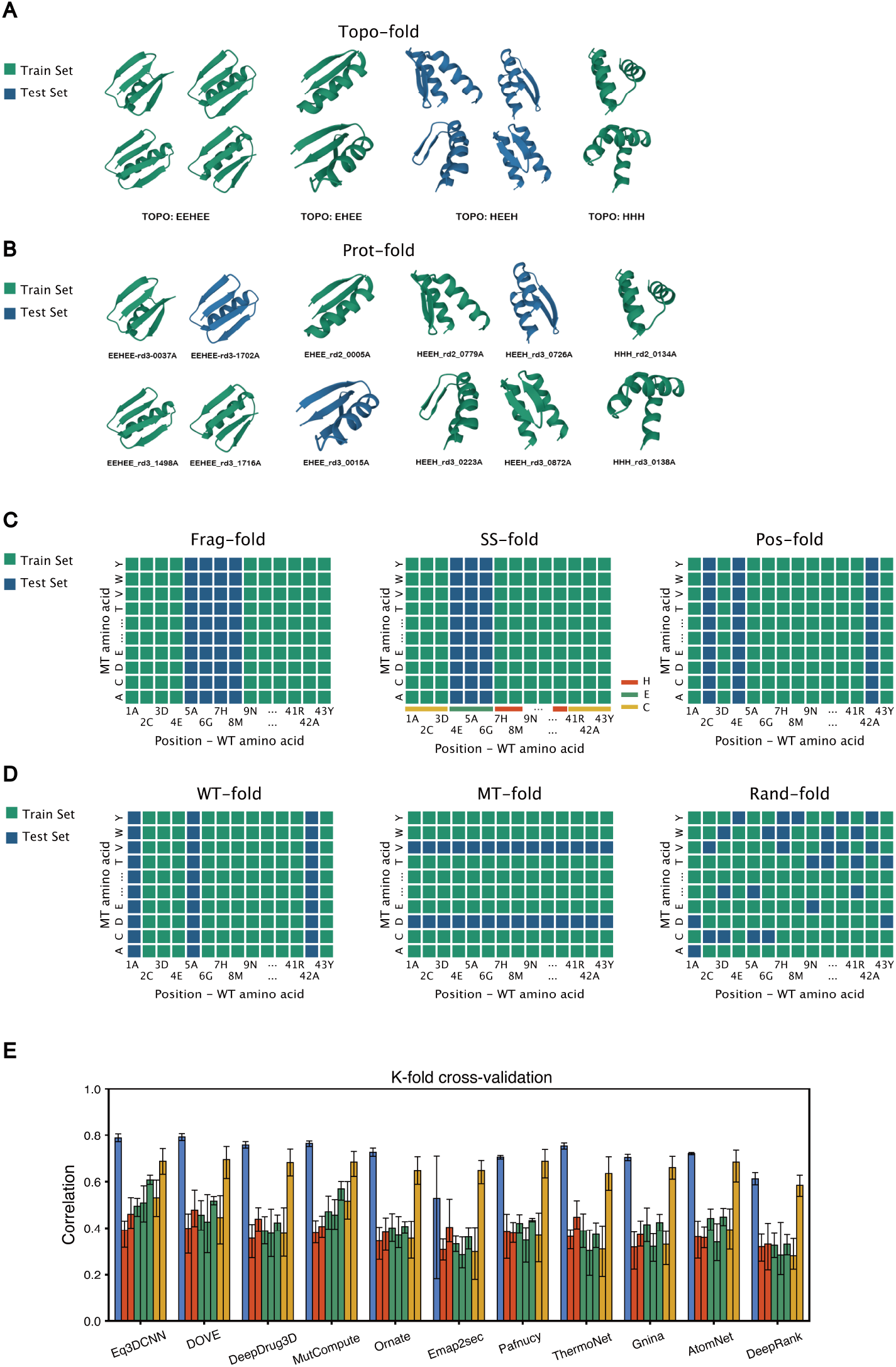
K-fold cross-validation setup and results. (**A**) Demonstration of the Topo-fold (k=4), that splits dataset by the topology. (**B**) Demonstration of the Prot-fold (k=5), that splits dataset by protein. (**C**) Demonstration of splitting dataset by positions. The Frag-fold (k=5) that splits dataset by a fragment (continuous positions), SS-fold (k=3) that splits dataset by the secondary structure (continuous positions within a secondary structure), and Pos-fold (k=5) that splits dataset by positions (random positions). (**D)** Demonstrations of splitting dataset by wildtype (WT-fold, k=5), mutation amino acid (MT-fold, k=5) and random splitting (Rand-fold, k=5), repectively. (**E)** Cross-validation results on Eq3DCNN and other methods.

## References

1. Sou Sugiki, Teppei Niide, Yoshihiro Toya, and Hiroshi Shimizu. Logistic regression-guided identification of cofactor specificity-contributing residues in enzyme with sequence datasets partitioned by catalytic properties. ACS Synthetic Biology, 11(12):3973–3985, 2022.

2. Yiquan Wang, Meng Yuan, Huibin Lv, Jian Peng, Ian A Wilson, and Nicholas C Wu. A large-scale systematic survey reveals recurring molecular features of public antibody responses to sars-cov-2. Immunity, 55(6):1105–1117, 2022.

3. Carlos L Araya, Douglas M Fowler, Wentao Chen, Ike Muniez, Jeffery W Kelly, and Stanley Fields. A fundamental protein property, thermodynamic stability, revealed solely from large-scale measurements of protein function. Proceedings of the National Academy of Sciences, 109(42):16858–16863, 2012.

4. Angela M Phillips, Katherine R Lawrence, Alief Moulana, Thomas Dupic, Jeffrey Chang, Milo S Johnson, Ivana Cvijovic, Thierry Mora, Aleksandra M Walczak, and Michael M Desai. Binding affinity landscapes constrain the evolution of broadly neutralizing anti-influenza antibodies. Elife, 10:e71393, 2021.

5. Kevin K Yang, Zachary Wu, and Frances H Arnold. Machine-learning-guided directed evolution for protein engineering. Nature methods, 16(8):687–694, 2019.

6. Patrick Koenig, Chingwei V Lee, Benjamin T Walters, Vasantharajan Janakiraman, Jeremy Stinson, Thomas W Patapoff, and Germaine Fuh. Mutational landscape of antibody variable domains reveals a switch modulating the interdomain conformational dynamics and antigen binding. Proceedings of the National Academy of Sciences, 114(4):E486–E495, 2017.

7. Brian L Hie, Varun R Shanker, Duo Xu, Theodora UJ Bruun, Payton A Weidenbacher, Shaogeng Tang, Wesley Wu, John E Pak, and Peter S Kim. Efficient evolution of human antibodies from general protein language models. Nature Biotechnology, 2023.

8. Zachary Wu, SB Jennifer Kan, Russell D Lewis, Bruce J Wittmann, and Frances H Arnold. Machine learning-assisted directed protein evolution with combinatorial libraries. Proceedings of the National Academy of Sciences, 116(18):8852–8858, 2019.

9. Tianhao Yu, Haiyang Cui, Jianan Canal Li, Yunan Luo, Guangde Jiang, and Huimin Zhao. Enzyme function prediction using contrastive learning. Science, 379(6639):1358–1363, 2023.

10. Chloe Hsu, Hunter Nisonoff, Clara Fannjiang, and Jennifer Listgarten. Learning protein fitness models from evolutionary and assay-labeled data. Nature biotechnology, 40(7):1114–1122, 2022.

11. Surojit Biswas, Grigory Khimulya, Ethan C Alley, Kevin M Esvelt, and George M Church. Low-n protein engineering with data-efficient deep learning. Nature methods, 18(4):389–396, 2021.

12. Jung-Eun Shin, Adam J Riesselman, Aaron W Kollasch, Conor McMahon, Elana Simon, Chris Sander, Aashish Manglik, Andrew C Kruse, and Debora S Marks. Protein design and variant prediction using autoregressive generative models. Nature communications, 12(1):2403, 2021.

13. John Jumper, Richard Evans, Alexander Pritzel, Tim Green, Michael Figurnov, Olaf Ronneberger, Kathryn Tunyasuvunakool, Russ Bates, Augustin Žídek, Anna Potapenko, Alex Bridgland, Clemens Meyer, Simon A A Kohl, Andrew J Ballard, Andrew Cowie, Bernardino Romera-Paredes, Stanislav Nikolov, Rishub Jain, Jonas Adler, Trevor Back, Stig Petersen, David Reiman, Ellen Clancy, Michal Zielinski, Martin Steinegger, Michalina Pacholska, Tamas Berghammer, Sebastian Bodenstein, David Silver, Oriol Vinyals, Andrew W Senior, Koray Kavukcuoglu, Pushmeet Kohli, and Demis Hassabis. Highly accurate protein structure prediction with AlphaFold. Nature, 596(7873):583–589, 2021.

14. Alexander Rives, Joshua Meier, Tom Sercu, Siddharth Goyal, Zeming Lin, Jason Liu, Demi Guo, Myle Ott, C Lawrence Zitnick, Jerry Ma, et al. Biological structure and function emerge from scaling unsupervised learning to 250 million protein sequences. Proceedings of the National Academy of Sciences, 118(15):e2016239118, 2021. bioRxiv 10.1101/622803.

15. Joshua Meier, Roshan Rao, Robert Verkuil, Jason Liu, Tom Sercu, and Alexander Rives. Language models enable zero-shot prediction of the effects of mutations on protein function. bioRxiv, 2021.

16. Zeming Lin, Halil Akin, Roshan Rao, Brian Hie, Zhongkai Zhu, Wenting Lu, Nikita Smetanin, Robert Verkuil, Ori Kabeli, Yaniv Shmueli, Allan dos Santos Costa, Maryam Fazel-Zarandi, Tom Sercu, Salvatore Candido, and Alexander Rives. Evolutionary-scale prediction of atomic-level protein structure with a language model. Science, 379(6637):1123–1130, 2023. Earlier versions as preprint: bioRxiv 2022.07.20.500902.

17. Wen Torng and Russ B Altman. 3d deep convolutional neural networks for amino acid environment similarity analysis. BMC bioinformatics, 18:1–23, 2017.

18. Izhar Wallach, Michael Dzamba, and Abraham Heifets. Atomnet: a deep convolutional neural network for bioactivity prediction in structure-based drug discovery. arXiv preprint arXiv:1510.02855, 2015.

19. Sai Raghavendra Maddhuri Venkata Subramaniya, Genki Terashi, and Daisuke Kihara. Protein secondary structure detection in intermediate-resolution cryo-em maps using deep learning. Nature methods, 16(9):911–917, 2019.

20. Limeng Pu, Rajiv Gandhi Govindaraj, Jeffrey Mitchell Lemoine, Hsiao-Chun Wu, and Michal Brylinski. Deepdrug3d: classification of ligand-binding pockets in proteins with a convolutional neural network. PLoS computational biology, 15(2):e1006718, 2019.

21. Xiao Wang, Genki Terashi, Charles W Christoffer, Mengmeng Zhu, and Daisuke Kihara. Protein docking model evaluation by 3d deep convolutional neural networks. Bioinformatics, 36(7):2113–2118, 2020.

22. Guillaume Pages, Benoit Charmettant, and Sergei Grudinin. Protein model quality assessment using 3d oriented convolutional neural networks. Bioinformatics, 35(18):3313–3319, 2019.

23. Bian Li, Yucheng T Yang, John A Capra, and Mark B Gerstein. Predicting changes in protein thermodynamic stability upon point mutation with deep 3d convolutional neural networks. PLoS computational biology, 16(11):e1008291, 2020.

24. Raghav Shroff, Austin W Cole, Daniel J Diaz, Barrett R Morrow, Isaac Donnell, Ankur Annapareddy, Jimmy Gollihar, Andrew D Ellington, and Ross Thyer. Discovery of novel gain-of-function mutations guided by structure-based deep learning. ACS synthetic biology, 9(11):2927–2935, 2020.

25. Anastasiya V Kulikova, Daniel J Diaz, James M Loy, Andrew D Ellington, and Claus O Wilke. Learning the local landscape of protein structures with convolutional neural networks. Journal of Biological Physics, 47(4):435–454, 2021.

26. Marta M Stepniewska-Dziubinska, Piotr Zielenkiewicz, and Pawel Siedlecki. Development and evaluation of a deep learning model for protein–ligand binding affinity prediction. Bioinformatics, 34(21):3666–3674, 2018.

27. Matthew Ragoza, Joshua Hochuli, Elisa Idrobo, Jocelyn Sunseri, and David Ryan Koes. Protein–ligand scoring with convolutional neural networks. Journal of chemical information and modeling, 57(4):942–957, 2017.

28. Nicolas Renaud, Cunliang Geng, Sonja Georgievska, Francesco Ambrosetti, Lars Ridder, Dario F Marzella, Manon F Reau, Alexandre MJJ Bonvin, and Li C Xue. Deeprank: a deep learning framework for data mining 3d protein-protein interfaces. Nature communications, 12(1):7068, 2021.

29. Alex Krizhevsky, Ilya Sutskever, and Geoffrey E Hinton. Imagenet classification with deep convolutional neural networks. Advances in neural information processing systems, 25, 2012.

30. Taco Cohen and Max Welling. Group equivariant convolutional networks. In International conference on machine learning, pages 2990–2999. PMLR, 2016.

31. Maurice Weiler and Gabriele Cesa. General e (2)-equivariant steerable cnns. Advances in neural information processing systems, 32, 2019.

32. Kaiming He, Xiangyu Zhang, Shaoqing Ren, and Jian Sun. Deep residual learning for image recognition. In Proceedings of the IEEE conference on computer vision and pattern recognition, pages 770–778, 2016.

33. MD Shaji Kumar, K Abdulla Bava, M Michael Gromiha, Ponraj Prabakaran, Koji Kitajima, Hatsuho Uedaira, and Akinori Sarai. Protherm and pronit: thermodynamic databases for proteins and protein–nucleic acid interactions. Nucleic acids research, 34(suppl 1):D204–D206, 2006.

34. Joicymara S Xavier, Thanh-Binh Nguyen, Malancha Karmarkar, Stephanie Portelli, Pamela M Rezende, Joao PL Velloso, David B Ascher, and Douglas EV Pires. Thermomutdb: a thermodynamic database for missense mutations. Nucleic acids research, 49(D1):D475–D479, 2021.

35. Jan Stourac, Juraj Dubrava, Milos Musil, Jana Horackova, Jiri Damborsky, Stanislav Mazurenko, and David Bednar. Fireprotdb: database of manually curated protein stability data. Nucleic acids research, 49(D1):D319–D324, 2021.

36. Anna Jarzab, Nils Kurzawa, Thomas Hopf, Matthias Moerch, Jana Zecha, Niels Leijten, Yangyang Bian, Eva Musiol, Melanie Maschberger, Gabriele Stoehr, et al. Meltome atlas—thermal proteome stability across the tree of life. Nature methods, 17(5):495–503, 2020.

37. Gabriel J Rocklin, Tamuka M Chidyausiku, Inna Goreshnik, Alex Ford, Scott Houliston, Alexander Lemak, Lauren Carter, Rashmi Ravichandran, Vikram K Mulligan, Aaron Chevalier, et al. Global analysis of protein folding using massively parallel design, synthesis, and testing. Science, 357(6347):168–175, 2017.

38. Renxiao Wang, Xueliang Fang, Yipin Lu, and Shaomeng Wang. The pdbbind database: Collection of binding affinities for protein-ligand complexes with known three-dimensional structures. Journal of medicinal chemistry, 47(12):2977–2980, 2004.

39. Inyup Paik, Phuoc HT Ngo, Raghav Shroff, Daniel J Diaz, Andre C Maranhao, David JF Walker, Sanchita Bhadra, and Andrew D Ellington. Improved bst dna polymerase variants derived via a machine learning approach. Biochemistry, 62(2):410–418, 2021.

40. Hongyuan Lu, Daniel J Diaz, Natalie J Czarnecki, Congzhi Zhu, Wantae Kim, Raghav Shroff, Daniel J Acosta, Bradley R Alexander, Hannah O Cole, Yan Zhang, et al. Machine learning-aided engineering of hydrolases for pet depolymerization. Nature, 604(7907):662–667, 2022.

41. Taco S Cohen and Max Welling. Steerable cnns. arXiv preprint arXiv:1612.08498, 2016.

42. Maurice Weiler, Mario Geiger, Max Welling, Wouter Boomsma, and Taco S Cohen. 3d steerable cnns: Learning rotationally equivariant features in volumetric data. Advances in Neural Information Processing Systems, 31, 2018.

43. Douglas M Fowler and Stanley Fields. Deep mutational scanning: a new style of protein science. Nature methods, 11(8):801–807, 2014.

44. Kenneth A Matreyek, Lea M Starita, Jason J Stephany, Beth Martin, Melissa A Chiasson, Vanessa E Gray, Martin Kircher, Arineh Khechaduri, Jennifer N Dines, Ronald J Hause, et al. Multiplex assessment of protein variant abundance by massively parallel sequencing. Nature genetics, 50(6):874–882, 2018.

45. Karen S Sarkisyan, Dmitry A Bolotin, Margarita V Meer, Dinara R Usmanova, Alexander S Mishin, George V Sharonov, Dmitry N Ivankov, Nina G Bozhanova, Mikhail S Baranov, Onuralp Soylemez, et al. Local fitness landscape of the green fluorescent protein. Nature, 533(7603):397–401, 2016.

46. Gabriele Cesa, Leon Lang, and Maurice Weiler. A program to build E(N)-equivariant steerable CNNs. In International Conference on Learning Representations, 2022.

47. S Doerr, MJ Harvey, Frank Noe, and GHTMD De Fabritiis. Htmd: high-throughput molecular dynamics for molecular discovery. Journal of chemical theory and computation, 12(4):1845–1852, 2016.

